# Single Nucleotide Polymorphism (SNP) and Antibody-based Cell Sorting (SNACS): A tool for demultiplexing single-cell DNA sequencing data

**DOI:** 10.1101/2024.02.07.579345

**Authors:** VE Kennedy, R Roy, CAC Peretz, A Koh, E Tran, CC Smith, AB Olshen

## Abstract

**Motivation:** Recently, single-cell DNA sequencing (scDNA-seq) and multi-modal profiling with the addition of cell-surface antibodies (scDAb-seq) have provided key insights into cancer heterogeneity.

Scaling these technologies across large patient cohorts, however, is cost and time prohibitive. Multiplexing, in which cells from unique patients are pooled into a single experiment, offers a possible solution. While multiplexing methods exist for scRNAseq, accurate demultiplexing in scDNAseq remains an unmet need.

**Results:** Here, we introduce SNACS: Single-Nucleotide Polymorphism (SNP) and Antibody-based Cell Sorting. SNACS relies on a combination of patient-level cell-surface identifiers and natural variation in genetic polymorphisms to demultiplex scDNAseq data. We demonstrated the performance of SNACS on a dataset consisting of multi-sample experiments from patients with leukemia where we knew truth from single-sample experiments from the same patients. Using SNACS, accuracy ranged from 0.948 – 0.991 vs 0.552 – 0.934 using demultiplexing methods from the single-cell literature.

**Availability Implementation:** SNACS is available at https://github.com/olshena/SNACS.

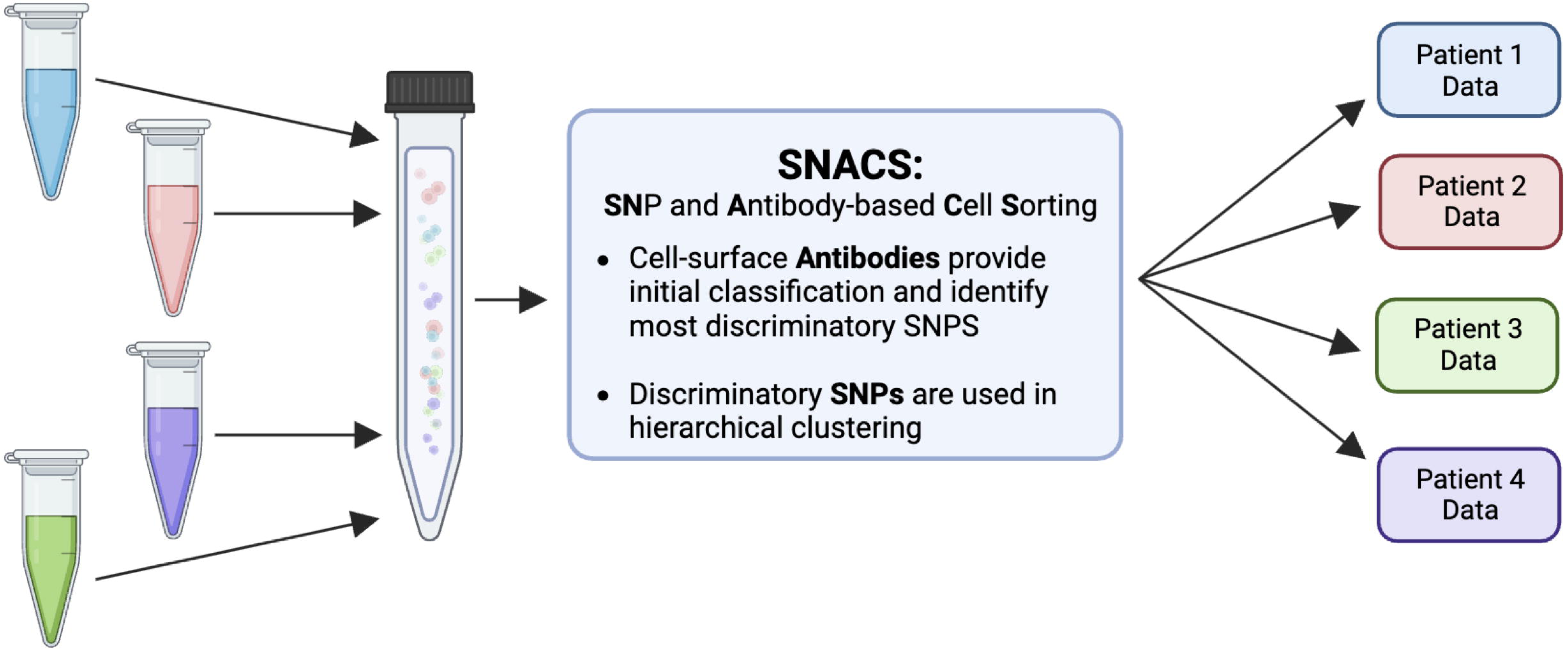

## 1 Introduction

Single-cell DNA sequencing (scDNA-seq) is an emerging microfluidic technology used in cancer research. Over the past several years, this technology has provided several key insights into cancer biology, intratumor heterogeneity, and clonal evolution^1,2^. Through direct measurement of mutational co-occurrence and acquisition, scDNA-seq can be used to reconstruct tumor phylogeny^3–7^, and serial measurements have further provided insight into treatment resistance and outcomes^8–10^. Through precise genetic profiling, scDNA-seq also provides improved ability to detect low-level disease and can thus distinguish clinically meaningful residual disease from non-cancerous populations^1,11^. More recently, scDNA-seq has been combined with single-cell measurements of cell-surface protein expression in a technology known as “scDAb-seq” for SC DNA and Antibody-seq^3,4,12^. This multi-omic technology provides novel insight into the complex relationship between cancer genotype and phenotype^3,4,12^. Taken together, scDNA-seq and scDAb-seq have opened a new frontier in cancer research.

Despite these abilities, there are several limitations to employing these technologies at scale. Both scDNA-seq and scDAb-seq are costly^1^ in terms of material and time needed to perform single cell assays, restricting adoption to highly-resourced research laboratories. To date, most scDNA-seq studies on human samples have included fewer than 10 patients and analysis of large patient cohorts and/or multiple timepoints per patient remains cost prohibitive. These costs limit the translation of single-cell technologies from research to viable clinical assays^13^.

One strategy for mitigating such challenges is *multiplexing*, in which cells from multiple unique individuals are pooled into a single microfluidic run and then subsequently *demutiplexed* using diverse bioinformatic tools. If employed successfully, this strategy can result in lower per sample library preparation costs and increased efficiency. Multiplexing, however, is highly error prone. In addition to incorrectly assigning cells to parent samples, multiplexing can result in *multiplets* where single cells from two or more individuals are encapsulated into a single droplet causing information from multiple individuals to be falsely associated with a single cell barcode. Without accurate identification and removal, multiplets may incorrectly appear to be unique cell populations and thus lead to inaccurate downstream analyses.

To date, single-cell multiplexing and multiplet identification has primarily been described in the single-cell RNAseq literature. Methods include barcode-based and single nucleotide polymorphism (SNP)-based approaches (**Figure 1A**). In barcode based approaches, cells from unique samples are labeled with sample-level DNA barcodes and attached either via cell-surface antibodies^14,15^, lipid-bound cell membrane tags^16^, or viral integration of DNA barcodes directly into the genome^17^. In SNP-based approaches, multiplexed samples are demultiplexed based on natural genetic polymorphisms or “endogenous” barcodes^18–21^. Both strategies have limitations. In barcode-based multiplexing, cells may be bound by multiple different sample-level barcodes, the incorrect barcode, or no barcode entirely. In SNP-based demultiplexing, multiplexed samples may be unclassifiable if sufficient discriminatory SNPs are not present or if sequencing depth is inadequate. Importantly, SNP-based demultiplexing is also dependent on *a priori* knowledge to assign cells to their sample of origin.

**Figure 1.**
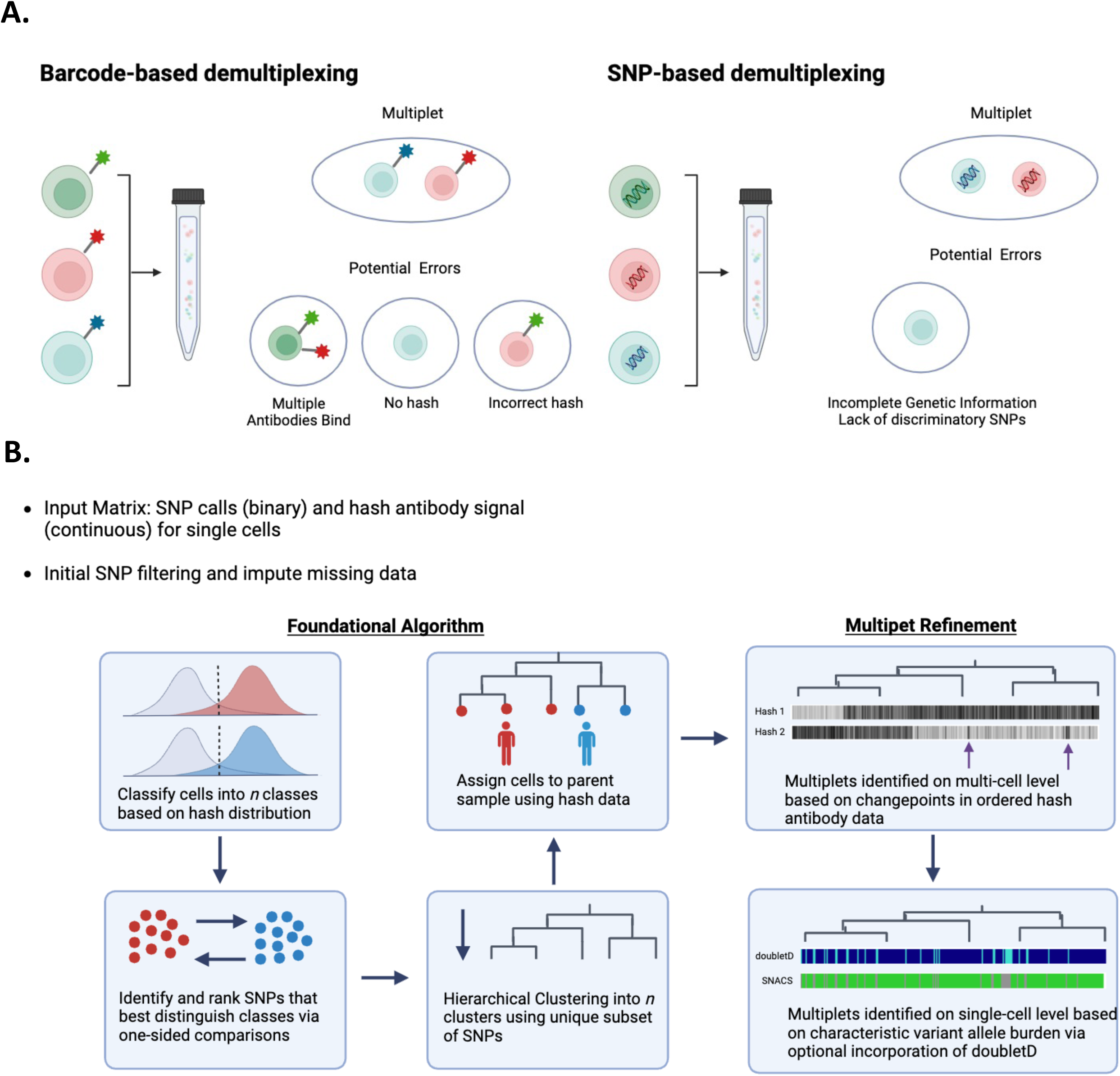
A. Existing demultiplexing approaches described in scRNA-seq include barcode- and SNP-based approaches. Both are imperfect and require identification of multiplets. B. Schematic of SNACS algorithm. SNACS offers a novel, combinatory approach using both SNPs and barcoded “hash” antibodies to demultiplex samples and resolve mutiplets.

In this work, we describe a novel scDNA-seq multiplexing approach, algorithm, and visualization methodology. Based upon DAbseq technology, this approach combines both SNP and barcoding based approaches for demultiplexing and multiplet identification, thus increasing accuracy. We have called this approach SNACS, for **SN**P and **A**ntibody-based **C**ell **S**orting. Our formulation is novel because previously, as far as we know, SNP and barcoding information have not been used in tandem for demultiplexing.

## 2 Algorithm

The SNACS algorithm described in detail here is outlined in **Figure 1B**. For each multiplexed experiment, SNPs are treated as binary (mutated or wildtype) and hash antibody expression is treated as continuous. SNP data are first filtered to remove both SNPs and single cells with high missingness. We used a threshold of 40% missing data for both, but results should not be sensitive to these choices. Hash antibody counts are normalized using the centered log ratio transformation, as is common in both SC DAbseq and CITEseq analysis^22^. In this normalization, the hash antibody count for every cell is divided by the geometric mean across cells and then log base 2 transformed.

### 2.1 Hash antibody data provides initial classification

First, hash antibody data is used to classify cells into *n* preliminary groups, where *n* is the number of parent samples and thus hash antibodies. For each hash antibody, the antibody expression for a multiplexed experiment is expected to be bimodal, with one right mode comprised of antibody-stained cells belonging to a single parent sample and one left mode comprised of unstained cells from alternate parent samples. To estimate the background antibody distribution, we generate a symmetric distribution by reflecting the data to the left of the left mode about that mode (**Figure S1).** Cells are assigned to a specific hash antibody if the antibody expression of that cell is expressed highly; we used a threshold of the 95^th^ percentile of the background distribution to estimate a cell as positive for a hash. Cells that were assigned to either multiple hashes or to no hashes were excluded in this step.

### 2.2 SNPs that separate preliminary groups are identified

Next, we select the SNPs that best distinguish the hash-antibody-defined groups from the initial preliminary classifications of the previous step. For each SNP, we compare the proportion of mutated (i.e., 1) values pairwise between hash groups and rank these comparisons using a one-sided chi-square test. Our test is one-sided to identify SNPs that have a higher proportion of positivity for each comparison. For each of *n* groups, *n – 1* comparisons are performed, and for each comparison, we choose the top *k* SNPs. The value of *k* is user-defined and we have set the default as three. The result of our comparisons identifies a total of *n*\*(*n – 1)*k* SNPs that separate groups, but as the same SNPs may be chosen for more than one comparison, we limit ourselves to the unique subset of SNPs.

### 2.3 SNPs are hierarchically clustered and hash antibody data refines those clusters and assigns them to parent samples

Next, we perform agglomerative hierarchical clustering of the binary SNP data from all cells using the unique subset of SNPs identified in the previous step. Prior to clustering, missing SNP data is imputed utilizing a majority vote of the five nearest neighbors from the kNN function of the VIM R Package. Clustering is performed using cosine as the distance function and Ward’s method for joining clusters. The resulting dendrogram is cut into *n* clusters. Clusters are further identified by traversing down the hierarchical tree and splitting if a significant difference is found when comparing the daughter nodes of any current node. The comparisons are made for every hash antibody, and the two-sample t-test with equal variances is used to test for differences with an unadjusted p-value threshold of 10^-5^. The process is stopped when no additional differences are found. We do not allow clusters of fewer than two cells.

Clusters are then assigned to a specific hash antibody and parent sample by comparing the antibody expression of the cluster to the hash background distributions as described in Section 2.1. In our analysis we assigned a cluster to a hash if > 50% of cells from that cluster have a hash expression that exceeded the 95^th^ percentile of the background distribution. Clusters assigned to multiple hashes are designated as multiplets, and clusters not assigned to a hash antibody are designated “no call”. In the SNACS R package, the output and visualization of this initial demultiplexing is designated “SNACS Round 1” (**Figures S2-4**).

### 2.3 Refined multiplet detection using Circular Binary Segmentation

As accurately detecting multiplets is a significant concern in multiplexed SC data, we perform up to two additional steps to improve multiplet detection. First, we refined the calling of multiplets at the multi-cell level using hash antibody data. To do this, we estimate the mean hash antibody expression for each hash based on the cell clusters that were uniquely assigned to that hash in SNACS Round 1 (Section 2.3). Then, we calculate the Euclidean distance to the cluster center for every cell and hash. Within each cluster, using the ordering of the cells determined by the clustering, we next segment the distance value for every hash by using circular binary segmentation (CBS), an algorithm designed to identify contiguous regions of homogenous DNA copy in the genome by estimating changepoints in sequential data^23^. The superset of all changepoints is then used to form new clusters, and those new clusters are then, possibly, reassigned in multiplets using the same approach as described in Section 2.3.

In this analysis, only narrow segments are considered; we chose 100 or fewer cells as narrow. To provide additional power in this step, our hash background cutoff was the 75th percentile instead of the 95^th^. In the SNACS R package, the output and visualization of this optional subsequent refinement step is designated “SNACS Round 2” (**Figures S2-4).**

### 2.4 Additional multiplet detection via combination with doubletD

We also include the capability to call multiplets at the single-cell level by incorporating the previously-published doubletD algorithm^24^. In doubletD, matrices of total and alternate allele depth for each single-cell barcode are used to identify doublets based on increased allele frequency and/or drop-out via an expectation-maximization approach. For each single cell in our multi-sample experiments, we considered the cell a multiplet if it was called a multiplet by SNACS, as detailed above, *or* by doubletD. In the SNACS R package, the output and visualization of this optional subsequent refinement step is designated “**SNACS plus doubletD”**. Of note, although the mathematics of the doubletD algorithm are identical to those previously published, we translated the author-supplied code from python to R for seamless incorporation into our software.

## 3 System and Methods

### 3.1 Generation of empiric data from leukemia patient samples

To evaluate the SNACS algorithm, we generated empiric data by conducting seven scDAb-seq experiments on combinations of pooled samples from four adult patients with acute myeloid leukemia using a microfluidic approach via the Tapestri platform (MissionBio). Briefly, cryopreserved cells were thawed, normalized to 10,000 cells/μL in 180 μL PBS (Corning), and then incubated with Human TruStain FcX (BioLegend) and salmon sperm DNA (Invitrogen) for 15 minutes at 4C. Cells were then incubated with oligo-conjugated cell-surface antibodies (TotalSeq “hashtag” antibodies, BioLegend) for 30 minutes to provide patient-level identifiers. Each patient was assigned a unique hash antibody, and patients were pooled together as described in Table 1.

**Table 1.** Experimental Plan for Generation of Empiric Single-Cell DNAseq Data.

| **Experiment** | **Patient** | **Hash** |
| --- | --- | --- |
| Experiment 1 | Patient A | TS-1 |
|  | Patient A | TS-2 |
| Experiment 2 | Patient B | TS-2 |
|  | Patient B | TS-3 |
| Experiment 3 | Patient C | TS-3 |
|  | Patient C | TS-4 |
| Experiment 4 | Patient D | TS-1 |
|  | Patient D | TS-4 |
| Experiment 5 | Patient A | TS-1 |
|  | Patient B | TS-2 |
| Experiment 6 | Patient B | TS-2 |
|  | Patient C | TS-3 |
|  | Patient D | TS-4 |
| Experiment 7 | Patient A | TS-1 |
|  | Patient B | TS-2 |
|  | Patient C | TS-3 |
|  | Patient D | TS-4 |

Next, pooled samples were resuspended in cell buffer (MissionBio), diluted to 4-7e6 cells/mL, and loaded onto a microfluidics cartridge, where individual cells were encapsulated, lysed, and barcoded using the Tapestri instrument. DNA from barcoded cells was amplified via PCR using a targeted panel that included 288 amplicons across 66 genes associated with acute myeloid leukemia (Supplementary Table 1). DNA PCR products were isolated, purified with AmpureXP beads (Beckman Coulter), used as a PCR template for library generation, and then repurified with AmpureXP beads. Protein PCR products from hash antibodies were isolated from the supernatant via incubation with a 5’ Biotin Oligo (ITD). Protein PCR products were then purified using Streptavidin C1 beads (Thermo Fisher Scientific), used as a PCR template for library generation, and then repurified using AmpureXP beads. Both DNA and protein libraries were quantified and assessed for quality via a Qubit fluorometer (Life Technologies) and Bioanalyzer (Agilent Technologies) prior to pooling for sequencing on an Illumina Novaseq.

FASTQ files were processed via an open-source pipeline as described previously^12^. Valid cell barcodes were called using the inflection point of the cell-rank plot in addition to the requirement that 60% of DNA intervals were covered by at least eight reads. Variants were called using GATK (v 4.1.3.0) according to GATK best practices^25^.

### 3.2 Estimation of accuracy

To test the accuracy of SNACS and other demultiplexing algorithms, we compared multi-sample experiments (Experiments 5-7) to single-sample experiments (Experiments 1-4). Our rationale was that the SNP profiles of the single-sample experiments would be reflected in the multi-sample experiments, thus allowing for an estimation of “truth”. Specifically, for each single cell in a multi-sample experiment, we assigned a “truth call” for each single cell by comparing the SNP profile of that cell against the SNP profile of the constituent single-sample experiments. We considered only SNPs that were genotyped in both the multi-sample and constituentsingle-sample experiments. **Supplemental Figures 5-7** provides visualization of accuracy calculations and assessments.

We determined truth calls as follows. For each single-sample experiment, if a SNP was mutated (i.e., 1) in >= 99.5% of cells, then it was considered to be positive. Similarly, if a SNP was wildtype (i.e., 0) in >= 99.5% of cells, then it was considered to be negative. A SNP was considered ambiguous if mutated in > 0.5% and < 99.5% of cells. Next, in the multi-sample experiment, we considered a single cell a true “singlet”, that is belonging to a specific single sample, when the SNP profile was an exact match for a constituent single-sample experiment, inconsistent with other single samples, and not a multiplet. A true multiplet was when the positive SNPs present were not possible from the positive or ambiguous SNPs present in any constituent single-sample experiment. An ambiguous call was made when a cell was not a singlet nor a multiplet.

For each of the three multi-sample experiments, we estimated total accuracy as the proportion of truth calls that were matched by the demultiplexing algorithm. We also came up with measures of sensitivity and specificity to characterize how well a demultiplexing algorithm identified multiplets. Sensitivity was defined as the proportion of true multiplets that were called multiplets and specificity as the proportion of true singlets that were called singlets. We additionally calculated the proportion of singlets assigned to an alternative single sample.

### 3.3 Comparison against Alternative Multiplexing Approaches

Using our benchmarking dataset, we evaluated SNACS against the following five diverse multiplexing approaches:

1. **HTOdemux** is a barcode-based approach from the Seurat package developed to demultiplex cells in scRNA-seq experiments based on k-medoid clustering of cell-surface hash antibody values^14^. We used this method with default parameters and without modifications.
2. **CiteFuse** is a barcode-based approach developed to demultiplex cells in scRNA-seq experiments using a Gaussian mixture model fit to log-transformed cell-surface hash antibody values^15^. We used this method with default parameters and without modifications.
3. **scSplit** is a SNP-based approach developed to demultiplex scRNA-seq data using initial k-means clustering followed by an expectation-maximization approach^19^. To modify this approach to scDNA-seq data, we used this method starting with an allele count matrix of SNPs x single-cell barcodes and then as-written without modifications.
4. While there are currently no named algorithms for demultiplexing scDNA-seq data, a SNP-based approach is described in a publication by **Robinson** et al^26^. This method uses initial k-means clustering followed by additional multiple identification by generating artificial multiplets and comparing them to true cells^26^. We refer to this method as the “Robinson method.”
5. **doubletD** is a SNP-based approach for detecting doublets in scDNA-seq data using an expectation-maximization approach, and is based on the observation the scDNA-seq multiplets tend to have an increase in number of allele copies and/or drop-out^24^. doubletD only detects whether a cell is a doublet vs not a doublet and does not demultiplex single cells or assign them to parent samples. Thus, in our comparison to doubletD, we compared only its ability to accurately identify a multiplet.

### 3.4 Implementation Details

SNACS software and associated documentation is freely available at https://github.com/olshena/SNACS. Version 1.0.3.4 was used in this analysis. Code used to generate truth calls and accuracy assessments is available as well at https://github.com/olshena/SNACS/tree/master/analysis. For experiments 1-7, both unprocessed FASTQ files as well as SNP and antibody matrices following FASTQ alignment and variant calling have been deposited in NCBI’s Gene Expression Omnibus (accession number GSE255224) and are available for public use.

## 4 Results

### 4.1 Performance of SNACS on Patient Data

We evaluated the performance of SNACS using empiric data from scDAb-seq of leukemia patients as outlined above. SNACS was able to accurately demultiplex multiplexed samples and identify multiplets (**Figure 2**; **Table 2; Tables S2-4**).

**Figure 2.**
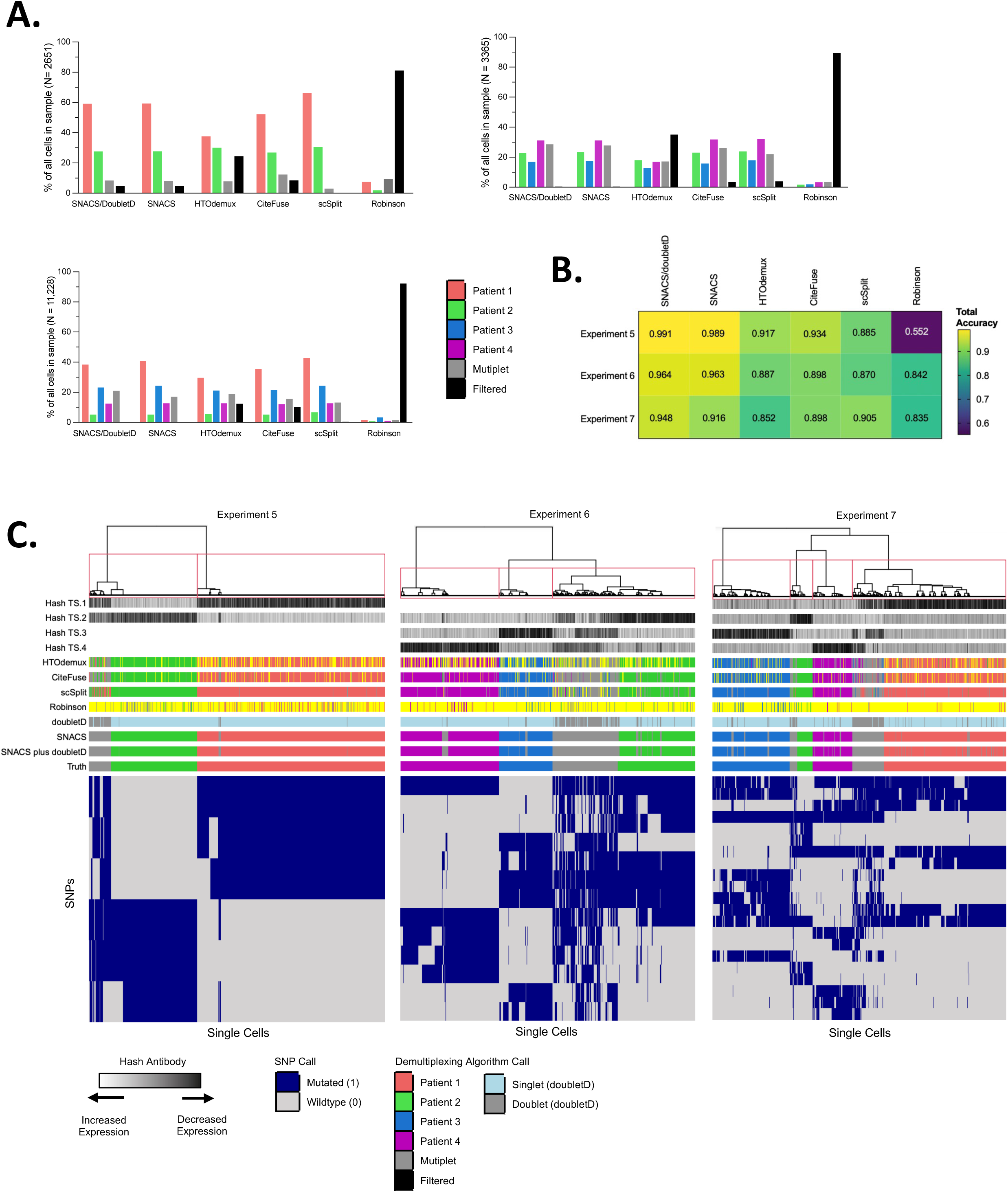
Comparison of SNACS to Alternate Demultiplexing Algorithms. A. Bar plots comparing relative proportions of singlets, multiplets, and filtered cells for SNACS plus alternative demultiplexing methods for multi-sample Experiment 5 (top left, patients 1 + 2), 6 (top right, patients 2 + 3 + 4), and 7 (bottom right, patients 1 + 2 + 3 +4). Relative to the comparison method s, SNACS filtered fewer cells. B. Heatmap of total accuracy for SNACS plus alternative demultiplexing methods (columns) for multi-sample Experiments 5-7 (rows). C. Heatmap of single cells (columns) vs SNPs used in final clustering by SNACS (rows) for multi-sample Experiments 5, 6, and 7. Rows at the top of the heatmap represent, in order from top to bottom, hash antibody signals; sample calls by HTOdemux, CiteFuse, scSplit, the Robinson method, doubletD, SNACS, SNACS plus doubletD, and truth calls.

**Table 2.** Accuracy Metrics for SNACS and other Demultiplexing Approaches.

| Algorithm | % Cells Filtered | Total Accuracy | Sensitivity | Specificity | No. of True Singlets | No. of True Multiplets | Fraction of Called Singlets which are True Alternative Singlets |
| --- | --- | --- | --- | --- | --- | --- | --- |
| **Experiment 5 (N = 2651 Cells)** |  |  |  |  |  |  |  |
| SNACS | 4.90% | **0.991** | 0.980 | **0.992** | 2320 | 200 | 0.000 |
| SNACS or doubletD | 4.90% | 0.989 | **0.985** | 0.990 | 2320 | 200 | 0.000 |
| HTOdemux | 24.44% | 0.917 | 0.581 | 0.979 | 1692 | 310 | 0.005 |
| CiteFuse | 8.49% | 0.934 | 0.822 | 0.971 | 2099 | 326 | 0.001 |
| scSplit | **0.15%** | 0.885 | 0.187 | 0.984 | 2319 | 327 | 0.008 |
| Robinson | 81.10% | 0.552 | 0.969 | 0.526 | 468 | 32 | 0.000 |
| doubletD | 0.0% | 0.978 | 0.942 | 0.983 | 2320 | 330 | N/A |
| **Experiment 6 (N = 3365 Cells)** |  |  |  |  |  |  |  |
| SNACS | 0.446% | **0.964** | 0.976 | **0.959** | 2484 | 837 | 0.0004 |
| SNACS or doubletD | 0.446% | 0.963 | **0.992** | 0.953 | 2484 | 837 | 0.0004 |
| HTOdemux | 35.00% | 0.887 | 0.758 | 0.905 | 1617 | 561 | 0.005 |
| CiteFuse | 3.45% | 0.898 | 0.828 | 0.922 | 2408 | 812 | 0.004 |
| scSplit | 3.95% | 0.870 | 0.719 | 0.916 | 2455 | 752 | 0.004 |
| Robinson | 89.48% | 0.842 | 0.790 | 0.863 | 249 | 105 | 0.000 |
| doubletD | 0.0% | 0.919 | 0.771 | 0.970 | 2485 | 849 | N/A |
| **Experiment 7 (N = 11,228 Cells)** |  |  |  |  |  |  |  |
| SNACS | 0.0% | **0.948** | 0.919 | **0.953** | 8517 | 1588 | 0.0002 |
| SNACS or doubletD | 0.0% | 0.916 | **0.948** | 0.910 | 8517 | 1588 | 0.0002 |
| HTOdemux | 12.31% | 0.852 | 0.759 | 0.872 | 7392 | 1558 | 0.015 |
| CiteFuse | 10.22% | 0.898 | 0.771 | 0.925 | 7557 | 1562 | 0.008 |
| scSplit | 0.39% | 0.905 | 0.677 | 0.948 | 8494 | 1573 | 0.009 |
| Robinson | 92.20% | 0.835 | 0.705 | 0.845 | 776 | 61 | 0.003 |
| doubletD | 0.0% | 0.917 | 0.876 | 0.923 | 8517 | 1588 | N/A |

Total Accuracy is defined as proportion correctly called cells/all called cells. Sensitivity is defined as proportion of True Multiplets that were called Multiplets. Specificity is defined as proportion of True Singlets that were called Singlets. Accuracy, Sensitivity, and Specificity are only calculated for cells that were both not filtered by the demultiplexing algorithm and non-ambiguous by truth call.

Experient 5 contained multiplexed samples from two individual patients. Of the 2,651 input cells, 2,521 (95.1%) had sufficient genotyping information to be included in the SNACS algorithm. Of the 70,476 unique input SNPs, SNACS identified a total of 6 unique SNPs that best defined the initial antibody-based classifications and were used in subsequent hierarchical clustering (**Table S5).** Of the 2,651 single cells, SNACS assigned 1,585 (59.8%) to Patient A, 735 (27.7%) to Patient B, and 200 (7.5%) as multiplets. When compared against the truth assessments, SNACS provided a total accuracy of 0.991, sensitivity (called multiplets/true multiplets) of 0.980, and specificity (called singlets/true singlets) of 0.992. With the addition of doubletD, sensitivity improved to 0.985, while specificity and total accuracy decreased to 0.990 and 0.981, respectively.

Experiment 6 contained multiplexed samples from three individual patients. Of the 3365 input cells, nearly all (99.6%) had sufficient genotyping information and were included in downstream analysis. SNACS assigned 799 (23.7%) to Patient A, 576 (17.1%) to Patient B, 1110 (33.0%) to Patient C, and 849 (25.2%) as multiplets. When compared against the truth assessments, SNACS provided a total accuracy of 0.964, sensitivity of 0.976, and specificity of 0.959. With the addition of doubletD, sensitivity again improved marginally to 0.992, while specificity decreased to 0.953.

Finally, Experiment 7 contained multiplexed samples from four individual patients. Of the 11,228 input cells, all (100%) had sufficient genotyping information and were included in downstream analysis. SNACS assigned 3,753 (33.4%), 490 (4.4%), 2,822 (25.1%), and 1,452 (12.9%) to Patients A, B, C and D, respectively; 1,588 (14.1%) were assigned as multiplets. SNACS provided a total accuracy of 0.948, sensitivity of 0.919, and specificity of 0.953. Like Experiments 5 and 6, the additional of doubletD improved sensitivity (0.948) while decreasing specificity (0.910).

Crucially, across all three multi-sample experiments, SNACS very rarely assigned singlets to incorrect parent samples, occurring in 0% of cells in Experiment 5, 0.04% of cells in Experiment 6, and 0.02% of cells in Experiment 7.

### 4.2 Comparison of SNACS vs Alternate Barcode- and SNP-based approaches

We also compared SNACS to 4 demultiplexing approaches and 1 doublet identification tool as outlined in Section 2.3. Relative to these comparison methods, SNACS preserved a high number of single cells, with < 5% of input cells filtered in Experiment 5 and <1% in Experiments 6 and 7 compared to 0.15% -81.1%, 3.5% - 89.5%, and 0.4% - 92.2% for the same experiments with the comparison methods (**Figure 2A**, **Table 2**). There was only once instance across methods and experiments (scSplit applied to Experiment 5) where an alternative method filtered fewer cells than SNACS.

SNACS also offered superior sensitivity and specificity in all experiments with sensitivities of 0.98, 0.98, and 0.92 relative to 0.19 – 0.97, 0.72 – 0.83, and 0.68 – 0.88 for Experiments 5, 6, and 7 for the comparison methods (**Table 2)**. Similarly, the specificity of SNACS was 0.99, 0.96, and 0.95 compared to 0.53 – 0.98, 0.86 – 0.92, and 0.85 – 0.93 for Experiments 5, 6, and 7 for the alternate methods (**Table 2)**. Finally, SNACS misidentified the lowest proportion of singlets as belonging to a true alternative parent sample across all 3 multi-sample experiments. After SNACS, there was no one demultiplexing approach that provided the best accuracy, sensitivity, and specificity metrics across all experiments. Interestingly, there was also not a consistent pattern of superior performance between SNP-based vs barcode-based algorithms **(Figure 2B)**.

Compared to the SNP-based methods scSplit and the Robinson method, SNACS employs minimal filtering, an approach chosen to both maximize the final number of assigned cells as well as to simplify our algorithm. SNACS does not filter for SNPs known to be common germline variants at the population level, but instead allows for unbiased selection for the most discriminatory SNPs. This differs from other SNP-based demultiplexing approaches, as scSplit first filters against SNPs of maximal variation detected via the 1000 Genomes Project^19^ and the Robinson method filters against SNPs extracted from NCBI dbSNP Build 144^26^.

When compared to the Robinson method, SNACS identified a relatively similar number of SNPs as contributing to final clustering with 7, 13, and 20 SNPs identified by SNACS vs 11, 5, and 16 SNPs identified by the Robinson method for Experiments 5, 6 and 7, respectively (**Table S5)**. By contrast, scType included 1142, 663, and 1620 SNPs as contributing to the initial clustering.

These SNPs had incomplete overlap, with SNACS and the Robinson method sharing 1, 3, and 5 SNPs and SNACS and scType sharing 5, 8, and 14 SNPs for Experiments 5, 6 and 7, respectively. Taken together, these data suggest that filtering SNP data through population-level databases may miss a large proportion of highly discriminatory SNPs.

## 5 Discussion

In recent years, the field of single-cell genomics has shifted from a handful of expert research laboratories to multiple research groups across diverse cancer histologies^27^. A robust demultiplexing approach provides one avenue for scaling and democratizing this emerging technology. While demultiplexing approaches have been developed for scRNA-seq, the robust translation of these methods to scDNA-seq remains an unmet need.

Here, we offer SNACS, a SNP-and-barcode-based algorithm for demultiplexing scDNA-seq data. SNACS is able to accurately assign singlets to parent samples without a priori knowledge of genetic features and, relative to existing approaches, provides greater sensitivity and specificity while simultaneously preserving a greater number of single cells. Finally, SNACS offers an accompanying data visualization, allowing the user to readily assess the quality of demultiplexing and decide whether additional optional multiplet refinement is appropriate for a particular dataset. While our experimental patient data was processed in an open source alignment pipeline^12^, SNACS is also compatible with output from the commercially available Mission Bio Tapestri Analysis system, although the SNP matrix must be formatted as binary (i.e., zygosity is not considered) and hash antibodies must first be normalized, such as by using the centered log ratio transformation.

It is clear from our investigations that there is demultiplexing information in data from both natural genetic variation and in patient-level hash antibodies. We chose to combine these two data types in a particular way, but other combinations might be effective as well. Specifically, we utilized the hash data to choose an initial discriminatory set of SNPs, we clustered those SNPs, and we used the hash data to further divide cells with similar genotypes. If more cells were available for each genotype, we would explore independently classifying each genotype by its hash distribution. Since we are limited, our classification borrows strength across similar genotypes.

Our experimental data was derived from a targeted DNA panel. This panel includes genomic markers of greatest interest in leukemia, similar to what is commonly used in biological and clinical investigation and was not designed to include SNPs with maximal variation. While this panel provided adequate SNP coverage for accurate demultiplexing, it is possible SNACS and other barcode-based approaches could be further optimized through rational design of DNA panels to include genomic regions of high population-level variation. A future panel could be designed to include SNPs previously identified in population studies as maximally variable^28,29^, or those commonly used in forensic analyses^30,31^. Y-chromosome encoded SNPs or short tandem repeat polymorphisms could also be included, and male and female patients could be intentionally multiplexed together^32,33^.

While SNACS provides many strengths in an area of unmet need, there are limitations to our approach that should be considered when it is used. Although accuracy was greater than against established methodologies, there were still a small proportion (<1.5%) of mis-classified singlets that were true multiplets. Should a biological experiment be conducted to identify small, rare cell populations, these mis-classified cells could result in false positive results. scDNA-seq experiments using SNACS will need to account for this, such as by discarding all cell populations smaller than a specific threshold. SNACS also requires that samples from different patients be multiplexed together. If an investigator wanted to analyze multiple timepoints from a single patient, samples from those timepoints would likely not be amenable to demultiplexing via SNACS.

Finally, SNACS is optimized to detect *across-sample* multiplets, in which multiplets are derived from different parent samples. In the single-cell literature, discussion also exists regarding *within-sample* multiplets, in which two or more cells from genotypically distinct subclones from the same parent sample are captured in a single droplet, resulting in a novel genotype derived from a single parent sample^15,24,34^. Unlike across-sample multiplets, within-sample multiplets may be present in single-sample experiments as well as multiplexed experiments. The true incidence of within-sample doublets in scDNAseq is unknown, and given their nature, they may not be detected by our truth-calling approach and algorithm. While SNACS does not directly detect within-sample multiplets, doubletD, which relies upon variant allele frequencies to detect multiplets, is optimized for both across- and within-sample doublets^24^, and it is possible the “SNACS plus doubletD” output identifies within-sample multiplets as well. Indeed, the decreased sensitivity of “SNACS plus doubletD” may in fact reflect the identification of within-sample multiplets not identified by our truth calling approach.

Questions also remain regarding the number of patients that could successfully be multiplexed together. While our approach found a slight decrease in sensitivity and specificity with an increase from two to four multiplexed samples, we do not know whether this decrease would continue with increasing sample numbers, and if so, at what rate, Similarly, while we did not observe an increase in proportion of multiplets with an increasing number of multiplexed samples, it is unknown whether multiplexing higher numbers of patients together would result in a greater proportion of multiplets. It may be that multiplexing is in fact limited by technical considerations. Assuming 20% multiplets per experiment, for an output of 7,500 cells, the median output from the Tapestri microfluidics platform used in our study, scaling beyond 6 patients would cause the number of singlets per patient to fall below 1,000; scaling beyond 60 would cause the number of singlets per patient to fall below 100. While the optimal number of cells per patient is largely dependent upon the scientific or clinical research question of interest and the importance of detecting rare populations, there are likely practical limitations to our approach.

Multiplexing scDNA-seq allows for efficient scaling of an emerging technology, and SNACS enables rapid and accurate demultiplexing and multiplet identification. Future studies are planned to validate and refine SNACS and evaluate it on larger numbers of multiplexed samples.

## Acknowledgements

We would like to acknowledge Aaron Logan for providing de-identified leukemia patient samples from the UCSF Hematologic Malignancies Tissue Bank. We would like to acknowledge Henrik Bengtsson for initial use and feedback on the SNACS R package. A.O. would like to acknowledge the UCSF Cancer Center Support Grant P30CA082103. C.S. is the Damon Runyon-Richard Lumsden Foundation Clinical Investigator supported (in part) by the Damon Runyon Cancer Research Foundation (CI-99-18) and is a Leukemia and Lymphoma Society Scholar in Clinical Research. Sequencing was performed at the UCSF CAT, supported by UCSF PBBR, RRP IMIA, and NIH 1S10OD028511-01 grants.

## References

1. Ediriwickrema, A., Gentles, A.J. & Majeti, R. Single-cell genomics in AML: extending the frontiers of AML research. Blood 141, 345–355 (2023).

2. Lim, B., Lin, Y. & Navin, N. Advancing Cancer Research and Medicine with Single-Cell Genomics. Cancer Cell 37, 456–470 (2020).

3. Miles, L.A., et al. Single-cell mutation analysis of clonal evolution in myeloid malignancies. Nature 587, 477–482 (2020).

4. Morita, K., et al. Clonal evolution of acute myeloid leukemia revealed by high-throughput single-cell genomics. Nat Commun 11, 5327 (2020).

5. Malikic, S., et al. PhISCS: a combinatorial approach for subperfect tumor phylogeny reconstruction via integrative use of single-cell and bulk sequencing data. Genome Res 29, 1860–1877 (2019).

6. Jahn, K., Kuipers, J. & Beerenwinkel, N. Tree inference for single-cell data. Genome Biol 17, 86 (2016).

7. Gawad, C., Koh, W. & Quake, S.R. Dissecting the clonal origins of childhood acute lymphoblastic leukemia by single-cell genomics. Proc Natl Acad Sci U S A 111, 17947–17952 (2014).

8. Peretz, C.A.C., et al. Single-cell DNA sequencing reveals complex mechanisms of resistance to quizartinib. Blood Adv 5, 1437–1441 (2021).

9. McMahon, C.M., et al. Clonal Selection with RAS Pathway Activation Mediates Secondary Clinical Resistance to Selective FLT3 Inhibition in Acute Myeloid Leukemia. Cancer Discov 9, 1050–1063 (2019).

10. Kennedy, V.E., et al. Multi-Omic Single-Cell Sequencing Reveals Genetic and Immunophenotypic Clonal Selection in Patients With FLT3-Mutated AML Treated With Gilteritinib/Venetoclax. Blood 140, 2244–2246 (2022).

11. Dillon, L.W., et al. Personalized Single-Cell Proteogenomics to Distinguish Acute Myeloid Leukemia from Non-Malignant Clonal Hematopoiesis. Blood Cancer Discov 2, 319–325 (2021).

12. Demaree, B., et al. Joint profiling of DNA and proteins in single cells to dissect genotype-phenotype associations in leukemia. Nat Commun 12, 1583 (2021).

13. Lei, Y., et al. Applications of single-cell sequencing in cancer research: progress and perspectives. J Hematol Oncol 14, 91 (2021).

14. Stoeckius, M., et al. Cell Hashing with barcoded antibodies enables multiplexing and doublet detection for single cell genomics. Genome Biol 19, 224 (2018).

15. Kim, H.J., Lin, Y., Geddes, T.A., Yang, J.Y.H. & Yang, P. CiteFuse enables multi-modal analysis of CITE-seq data. Bioinformatics 36, 4137–4143 (2020).

16. McGinnis, C.S., et al. MULTI-seq: sample multiplexing for single-cell RNA sequencing using lipid-tagged indices. Nat Methods 16, 619–626 (2019).

17. Guo, C., et al. CellTag Indexing: genetic barcode-based sample multiplexing for single-cell genomics. Genome Biol 20, 90 (2019).

18. Heaton, H., et al. Souporcell: robust clustering of single-cell RNA-seq data by genotype without reference genotypes. Nat Methods 17, 615–620 (2020).

19. Xu, J., et al. Genotype-free demultiplexing of pooled single-cell RNA-seq. Genome Biol 20, 290 (2019).

20. Huang, Y., McCarthy, D.J. & Stegle, O. Vireo: Bayesian demultiplexing of pooled single-cell RNA-seq data without genotype reference. Genome Biol 20, 273 (2019).

21. Kang, H.M., et al. Multiplexed droplet single-cell RNA-sequencing using natural genetic variation. Nat Biotechnol 36, 89–94 (2018).

22. Mule, M.P., Martins, A.J. & Tsang, J.S. Normalizing and denoising protein expression data from droplet-based single cell profiling. Nat Commun 13, 2099 (2022).

23. Olshen, A.B., Venkatraman, E.S., Lucito, R. & Wigler, M. Circular binary segmentation for the analysis of array-based DNA copy number data. Biostatistics 5, 557–572 (2004).

24. Weber, L.L., Sashittal, P. & El-Kebir, M. doubletD: detecting doublets in single-cell DNA sequencing data. Bioinformatics 37, i214–i221 (2021).

25. DePristo, M.A., et al. A framework for variation discovery and genotyping using next-generation DNA sequencing data. Nat Genet 43, 491–498 (2011).

26. Robinson, T.M., et al. Single-cell genotypic and phenotypic analysis of measurable residual disease in acute myeloid leukemia. Sci Adv 9, eadg0488 (2023).

27. Tanay, A. & Regev, A. Scaling single-cell genomics from phenomenology to mechanism. Nature 541, 331–338 (2017).

28. Yousefi, S., et al. A SNP panel for identification of DNA and RNA specimens. BMC Genomics 19, 90 (2018).

29. Kayser, M. & de Knijff, P. Improving human forensics through advances in genetics, genomics and molecular biology. Nat Rev Genet 12, 179–192 (2011).

30. Tillmar, A., Sturk-Andreaggi, K., Daniels-Higginbotham, J., Thomas, J.T. & Marshall, C. The FORCE Panel: An All-in-One SNP Marker Set for Confirming Investigative Genetic Genealogy Leads and for General Forensic Applications. Genes (Basel) 12(2021).

31. Budowle, B. & van Daal, A. Forensically relevant SNP classes. Biotechniques 44, 603–608, 610 (2008).

32. Jobling, M.A. & Tyler-Smith, C. Human Y-chromosome variation in the genome-sequencing era. Nat Rev Genet 18, 485–497 (2017).

33. Kayser, M. Forensic use of Y-chromosome DNA: a general overview. Hum Genet 136, 621–635 (2017).

34. Pellegrino, M., et al. High-throughput single-cell DNA sequencing of acute myeloid leukemia tumors with droplet microfluidics. Genome Res 28, 1345–1352 (2018).

