## Supplementary Figures for "Single Nucleotide Polymorphism (SNP) and Antibody-based Cell Sorting (SNACS): A tool for demultiplexing single-cell DNA sequencing data"

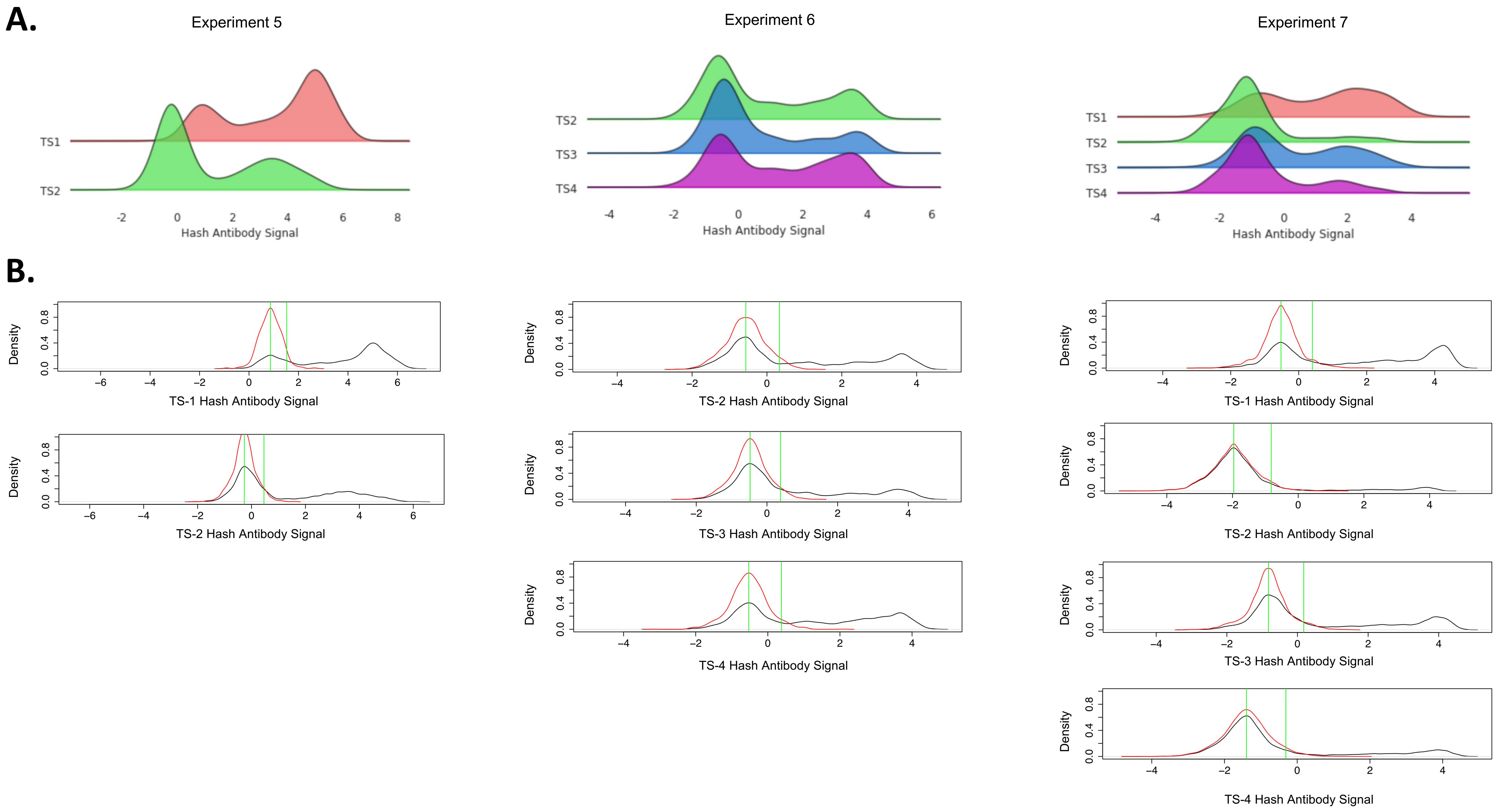


**Supplemental Figure 1. Hash Antibody Distribution for Multi-Sample Experiments**

A. Ridge plot of hash antibody expression for Experiment 5 (Patients A and B multiplexed), Experiment 6 (Patients B, C, and D multiplexed) and Experiment 7 (Patients 1, 2, 3, and 4 multiplexed.

B. In the foundational SNACS algorithm, SNP-defined clusters are assigned to a specific hash by comparing the actual antibody expression of the cluster (black line) to the expected hash antibody distribution. To generate the expected bimodal hash antibody distribution, we fit a Gaussian distribution to the left-most actual distribution (red line) and reflect the data to the left of the mode about the mode.

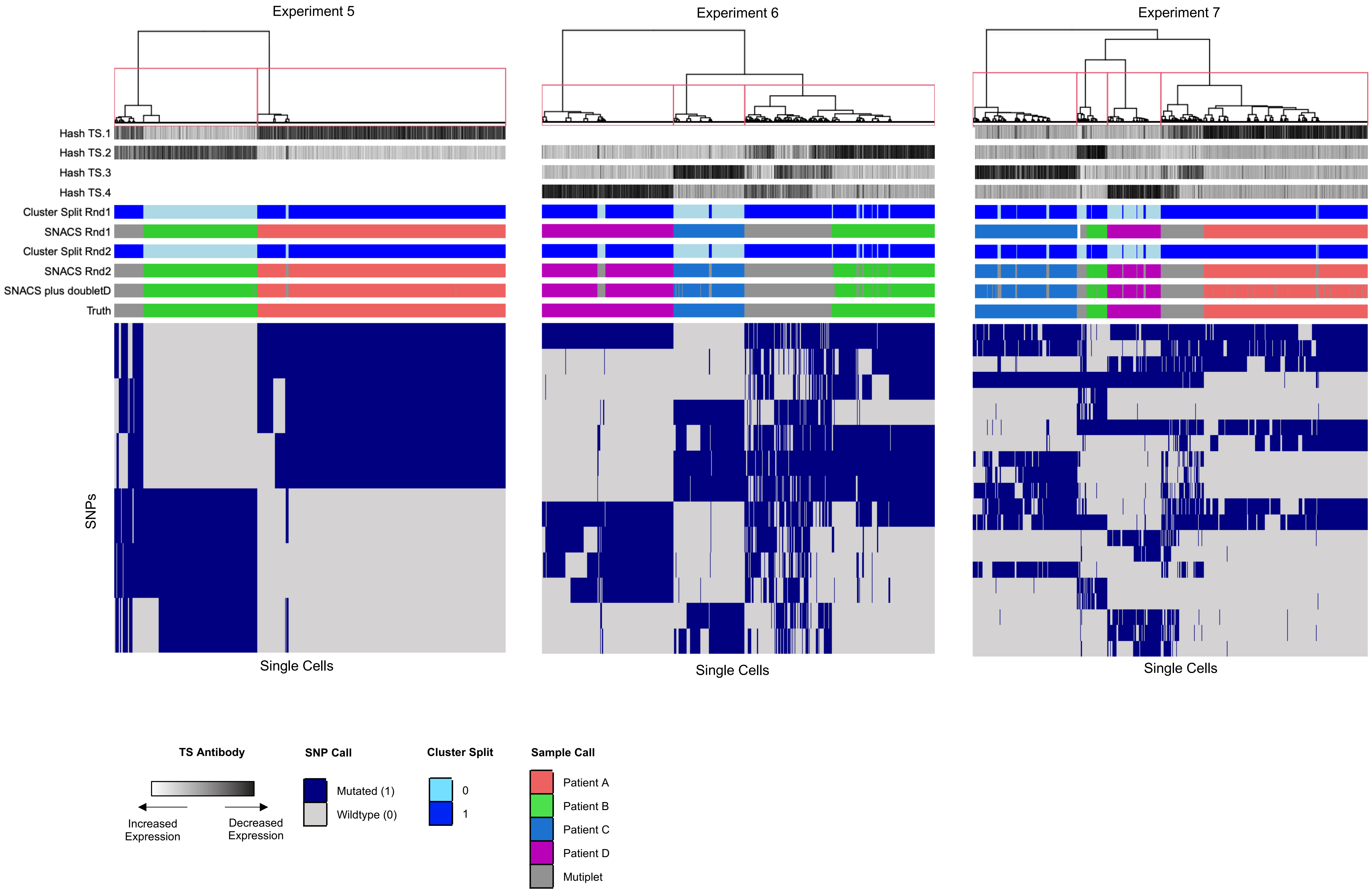


**Supplementary Figure 2. SNACS offers simple visualization of demultiplexing algorithm as shown by heatmaps of multi-sample Experiment 5 (Patient A and B), Experiment 6 (Patient B, C, and D) and Experiment 7 (Patient A, B, C, and D.)**
The heatmaps represents single cells (columns) vs SNPs (rows), color-coded by SNP mutational status. Rows above the heatmap represent, from top to bottom: hash antibody signal, cluster split and sample assignment from the foundational SNACs algorithm (SNACS Round 1), and cluster split and sample assignment from multiplet-refinement based on hash antibody signal (SNACS Round 2) and the inclusion of doubletD.

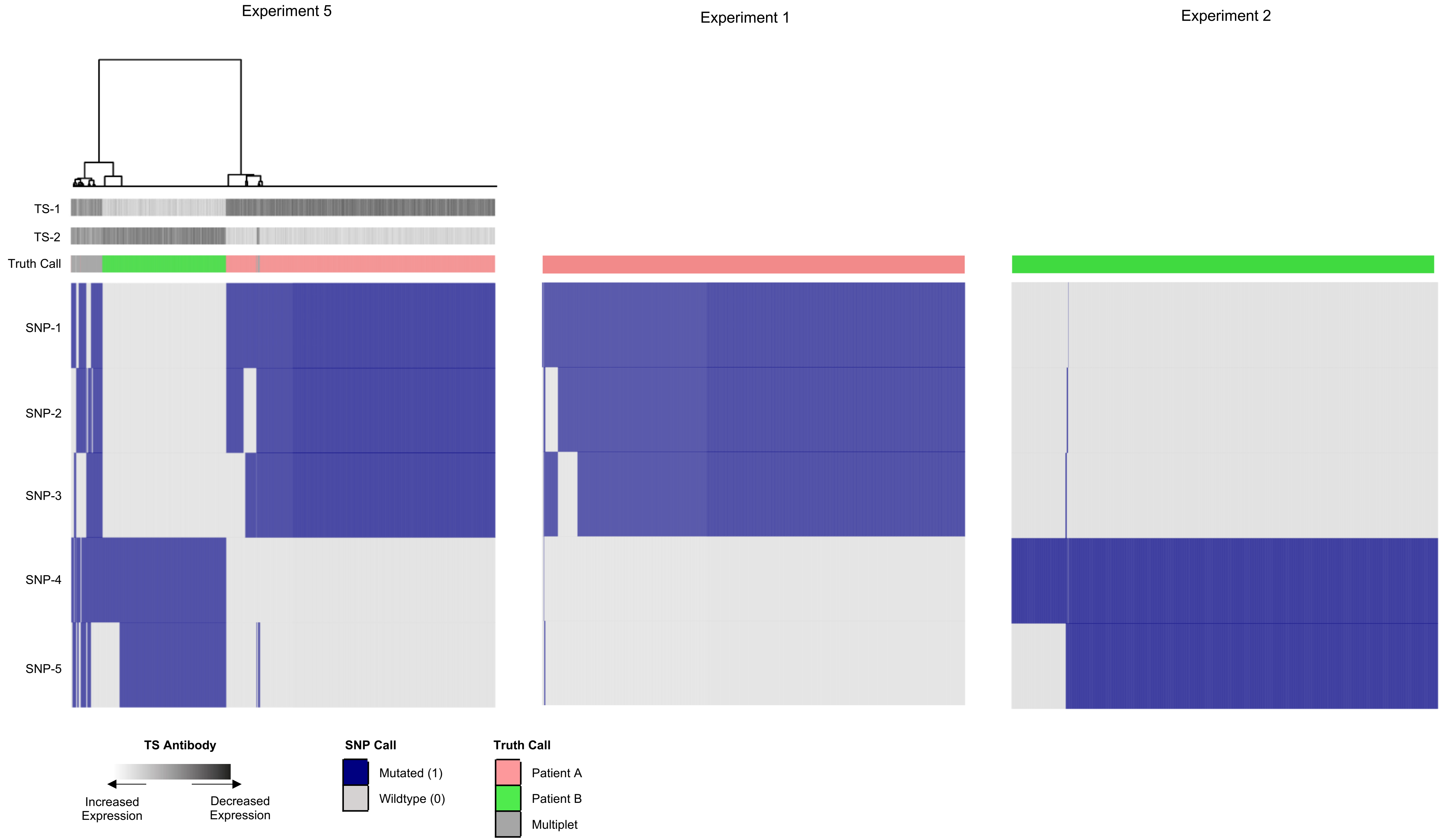


**Supplemental Figure 3. Heatmaps of multi-sample Experiment 5 (Patients A and B Mutliplexed) and single-sample Experiments 1 and 2 (Patients A and B separately) provide visualization of accuracy assessment and calculation.** The heatmap represents single cells (columns) vs SNPs (rows), color-coded by SNP mutational status. The 5 SNPs visualized are the SNPs which contributed to the final clustering in Experiment 5 which were also genotyped in Experiments 1 and 2. Rows above the heatmap represent, from top to bottom: hash antibody signal for TS-1 and TS2 and truth call as identified by single cell experiments.

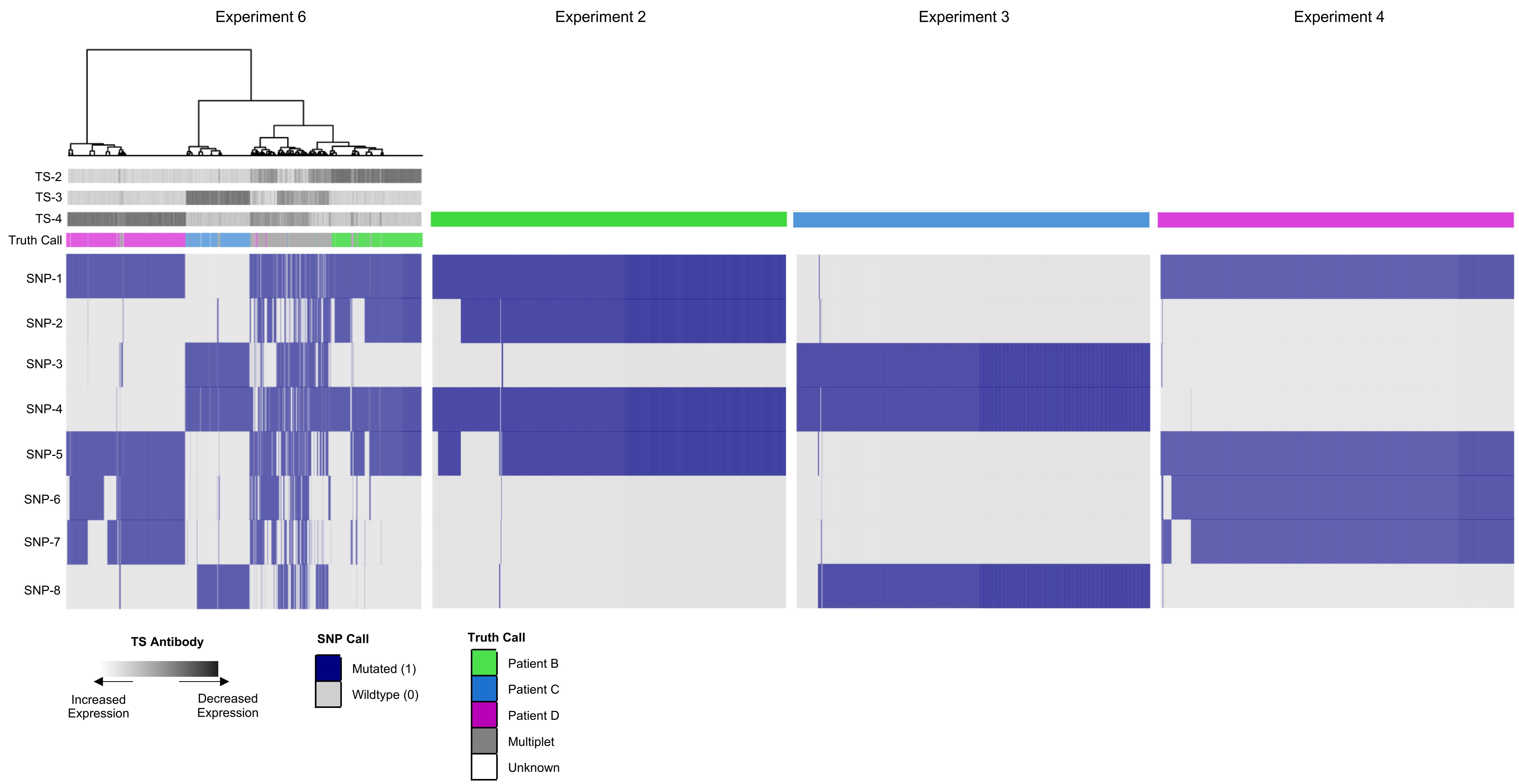


**Supplemental Figure 4. Heatmaps of multi-sample Experiment 6 (Patients B, C, and D Mutliplexed) and single-sample Experiments 2, 3, and 4 (Patients B, C, and D, respectively) provide visualization of accuracy assessment.** The heatmap represents single cells (columns) vs SNPs (rows), color-coded by SNP mutational status. The 7 SNPs visualized are the SNPs which contributed to the final clustering in Experiment 6 which were also genotyped in Experiments 2-4. Rows above the heatmap represent, from top to bottom: Hash antibody signal for TS-2, TS-3, and TS-4 and Truth Call as identified by single cell experiments.

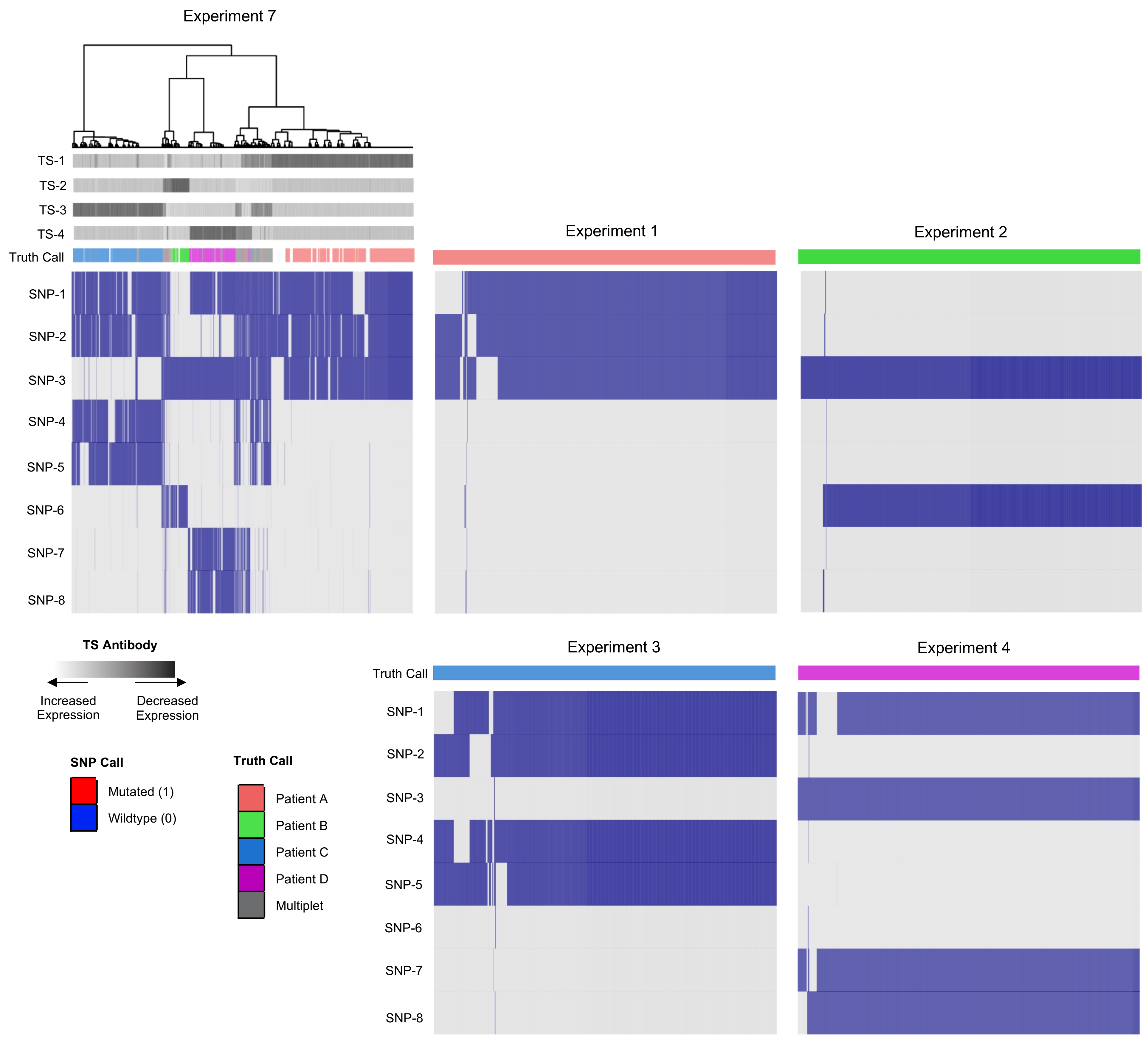


**Supplemental Figure 5. Heatmaps of multi-sample Experiment 7 (Patients A, B, C, and D Mutliplexed) and single-sample Experiments 1, 2, 3, and 4 (Patients A, B, C, and D, respectively) provide visualization of accuracy assessment.** The heatmap represents single cells (columns) vs SNPs (rows), color-coded by SNP mutational status. The 14 SNPs visualized are the SNPs which contributed to the final clustering in Experiment 7 which were also genotyped in Experiments 1-4. Rows above the heatmap represent, from top to bottom: Hash antibody signal for TS-1, TS-2, TS-3, and TS-4 and Truth Call as identified by single cell experiments.
